# Choices in Vaccine Trial Design for Epidemics of Emerging Infections

**DOI:** 10.1101/259606

**Authors:** Rebecca Kahn, Annette Rid, Peter G Smith, Nir Eyal, Marc Lipsitch

## Abstract

The 2014–2016 Ebola epidemic highlighted the lack of consensus on the design of trials for investigational vaccine products in an emergency setting. With the advent of the ring vaccination strategy, it also underscored that the range of design options is evolving according to scientific need and creativity. Ideally, principles and protocols will be drawn up in advance, facilitating expediency and trust, for rapid deployment early in an epidemic. Here, we attempt a summary of the scientific, ethical and feasibility considerations relevant to different trial designs. We focus on four elements of design choices which, in our view, are most fundamental to designing an experimental vaccine trial and for which the most distinctive issues arise in the setting of an emerging infectious disease for which no proven vaccines exist: 1) randomization unit, 2) trial population, 3) comparator intervention and 4) trial implementation. Likewise, we focus on three of several ethical considerations in clinical research, namely the trial’s social and scientific value, its risk-benefit profile and its participant selection. A catalogue of possible designs to guide trial design choices is offered, along with a systematic evaluation of the benefits and drawbacks of each in given contexts.

## Introduction

In outbreaks of emerging infectious diseases for which no proven efficacious vaccines exist, it is important both to test investigational vaccine products that passed safety and immunogenicity testing and then, if found efficacious, to deploy them rapidly for epidemic control. Following the 2014–2016 Ebola epidemic, the World Health Organization, the Coalition for Epidemic Preparedness Innovations, and other bodies committed to developing investigational vaccines for emerging infectious diseases.^1,2^ Although it may not be possible to assess the efficacy of these products in humans in advance of an epidemic, these organizations aim to evaluate their safety and immunogenicity, so that in the event of a future outbreak, they would be ready for efficacy testing and possible deployment. We use *efficacy* in this paper as shorthand *for protection against becoming infected* without prejudging various aspects of this protection which are called in various places efficacy and effectiveness.

The Ebola epidemic highlighted the lack of consensus on the design of efficacy trials for investigational vaccine candidates, and various strategies were deployed.^3^ Some argued for individually-randomized controlled trials (iRCTs), while others argued for forms of cluster-randomized trials (cRCTs).^4,5^ Notably, changes had to be made to some trial designs because of the rapidly declining disease incidence at, and following, trial implementation. Ideally, principles and protocols based on ethical, scientific and feasibility considerations will be drawn up in advance of an epidemic, facilitating expediency and trust, for rapid, early deployment. Here, we attempt a summary of the scientific, ethical and feasibility considerations relevant to different designs of Phase 3 vaccine trials. We lay out the possibilities for different trial designs to be considered for optimal vaccine testing during outbreaks of emerging infections. A catalogue of possible designs to guide trial design choices is offered, along with a systematic evaluation of the benefits and drawbacks of each in given contexts.

## Scope

When designing and implementing clinical trials of vaccines, important decisions must be made at different time points. Here we focus on key choices that must be made before efficacy trials (i.e. Phase 3 trials) during epidemics can begin, after initial safety and immunogenicity data (in Phase 1 and 2 trials) on the given investigational vaccine have been collected. There is a large literature on the use of observational, non-randomized designs to evaluate vaccine effectiveness, once a vaccine has been deployed in a control programme.^6–8^ However, we focus only on experimental, randomized trials in which there is some form of contemporaneous control group, as there is widespread consensus that such trials are the preferred option for assessing the efficacy of new vaccines, as they avoid the biases inherent in non-randomized comparisons. In the current regulatory system, experimental trials are generally considered the gold standard for vaccine licensure, and except in rare circumstances, they have been the required option for vaccine licensure.^9,10^ We also restrict our scope to settings in which no proven effective vaccine exists and to evaluation of a single vaccine;^11^ we do not discuss comparing more than one investigational vaccine in the same trial.

As part of the experimental design, choices must be made on the exact method of randomization – either at the individual or cluster level. A second choice pertains to the trial population. A third, to what intervention, if any, the comparison group will receive. A fourth one is of the strategy for implementing the trial. We thus discuss four key elements about vaccine trial design during epidemics: 1) randomization unit, 2) trial population, 3) comparator intervention and 4) trial implementation. These four elements of design choices are those which, in our view, are most fundamental to designing an experimental vaccine trial and raise the most distinctive dilemmas during outbreaks of emerging infections for which no proven vaccines exist. Likewise, we focus on three of several ethical considerations in clinical research, namely the trial’s social and scientific value, its risk-benefit profile and its participant selection.^12,13^

For each element, we discuss the advantages and disadvantages of possible design options in a single section that includes ethical, scientific and feasibility considerations, for all are interrelated. For example, social value and scientific validity are key ethical considerations for clinical research, as exposing participants to risks is justified only when the scientific methods are appropriate for answering the given research questions.^13,14^ Similarly, feasibility considerations have both ethical and scientific import because the value of trials cannot be realized without accounting for possible obstacles in their implementation.

## Randomized vaccine trial design choices during epidemics

Table 1 summarizes the major designs that have, to our knowledge, been used or proposed for vaccine trials. Some have never been employed in emergency settings but could be considered for future use, depending on the nature of the epidemic, including the mode of transmission of the epidemic agent. We reference the examples listed in the table as we discuss the four key elements in Sections I–IV.

**Table 1.** Trial design overview and examples of key trial designs to study efficacy/effectiveness of investigational vaccines. The table does not provide an exhaustive list of possible trial designs, in the sense that it does not include all logically possible combinations of options for the four choices we emphasize. Moreover, the innovation of the ring vaccination trial design during the 2014–6 Ebola epidemic shows that the range of design options is evolving according to scientific need and creativity.

| 1 | Comparison | 2 | Population | 3 | Implementation |
| --- | --- | --- | --- | --- | --- |
| A | Placebo | A | General | A | Parallel |
| B | Other / active | B | High-risk | B | Stepped |
| C | Delay or no vaccine |  |  | C | Ring (stepped) |

| # | Common name for trial design | 1 | 2 | 3 | Examples |
| --- | --- | --- | --- | --- | --- |
| **I. Individually randomized** |  |  |  |  |  |
| 1 | "Classic" individually-randomized controlled trial* | A B | A B | A | Pneumococcal vaccine trial, California<sup>55</sup><br>Rotavirus vaccine trial, Niger<sup>56</sup><br>PREVAIL Ebola vaccine trial, Liberia<sup>34</sup> |
| 2 | Sero-discordant couples design** | A | B | A | Herpes simplex virus, type 2 trial<sup>35</sup> |
| 3 | Individually randomized comparison to delayed vaccine | C | B | B | Ebola STRIVE trial, Sierra Leone, as performed<sup>36†</sup> |
| 4 | Individually-randomized controlled design with deliberately stepped rollout | A | A B | B | Proposed for Ebola vaccines<sup>49</sup> |
| **II. Cluster randomized** |  |  |  |  |  |
| 5 | "Classic" parallel, cluster-randomized controlled trial | A B | A | A | Pneumococcal vaccine trial, Navajo<sup>57</sup> |
| 6 | Stepped-wedge cluster-randomized design | C | A | B | Hepatitis B vaccine trial, Gambia<sup>48</sup><br>Ebola STRIVE trial, as initially proposed<sup>19</sup> |
| 7 | Cluster-randomized ring-vaccination trial vs. delayed vaccination | C | B | C | Ebola ça Suffit trial, Guinea<sup>15</sup> |

\* Most common design for vaccine efficacy trials. (A search on clinicaltrials.gov with filters "Interventional (or Clinical Trial)" for study type, "Phase 3" for study phase and search term "vaccine" for intervention resulted in 1251 trials, of which 989 were randomized. Out of a randomly selected 50 of these trials, all were individually randomized and 44 stated use of a parallel rollout.)
\*\* Seronegative partner of a seropositive person is at high-risk for exposure to infection and is randomized to vaccine or placebo.
† Unusual choice of delayed vaccination comparison due to perceived challenges of placebo in this setting.

### I. Randomization Unit

#### A. Overview

In efficacy trials of experimental vaccines, participants are randomized either at the individual level or in groups (clusters). By definition, in an iRCT, each individual is independently assigned at random to either the investigational vaccine or the control arm. The ratio of participants in the investigational vaccine arm to the control arm is typically 1:1 (i.e. vaccine:control), as this ratio is the most statistically efficient for obtaining a vaccine efficacy estimate with a fixed number of participants; however, a 2:1 or 3:1 ratio is sometimes used, e.g. when the investigators want to collect more data on the safety of the investigational vaccine. In a cRCT, clusters or groups of participants are randomized to the investigational vaccine or control arms. Often, the clusters are defined geographically (e.g. villages), but they can also be defined based on social contacts (e.g. Ebola ring vaccination trial).^15^

#### B. Features of individual randomization

A major advantage of individual randomization is its greater statistical efficiency compared to cluster randomization, meaning that iRCTs yield safety and efficacy estimates with a smaller confidence interval than cRCTs involving the same number of participants.^16^ In cRCTs, outcomes among members of the same cluster tend to be correlated with one another for reasons separate from the intervention received by that cluster. For example, they may share similar exposures or risk factors. A statistical “penalty”, known as the design effect, is therefore incurred when evaluating the uncertainty in the trial to account for such correlations.^17,18^ The design effect can be especially large when the incidence of a disease is highly clustered in space and time, as is often the case in emerging infectious diseases.^19–22^

However, it is important to note that iRCTs will generally only measure the *<u>direct</u>* protective effect of the investigational vaccine—that is, the extent by which the risk of disease is reduced when an individual is exposed to the infectious agent. The direct effect of a vaccine is mainly a characteristic of the vaccine itself and how it interacts with individuals, rather than of the way the vaccine is deployed in a particular population. For example, estimates of vaccine efficacy made in one population are routinely used in modelling exercises to estimate effectiveness or cost-effectiveness of the same vaccine in other populations, including modelling the indirect effect of the vaccine depending on vaccine coverage of the at-risk population. While direct vaccine effects are not always generalizable,^23–26^ they are often assumed to be “transportable” between settings because direct effects are measured at the individual level and not typically thought to depend much on the patterns of exposure and transmission in the population.

#### C. Features of cluster randomization

For some vaccines, specifically those for which there is person-to-person transmission of the infectious agent (including vector-borne or via an environmental reservoir), *<u>indirect</u>* effects may be an important component of their protective effect, when used in control programs. Such effects occur when the number of persons vaccinated in a community reduces the overall transmission rate of the infection in the community. This indirect effect (sometimes called *<u>herd</u>* effect) benefits both vaccinated and unvaccinated individuals, and the size of the effect will depend both on the level of the direct effect and the proportion of persons vaccinated in the population.^27^

In cRCT designs, the goal is to achieve high coverage of the investigational vaccine in some clusters, while those in other clusters do not receive the vaccine and serve as contemporaneous controls. If transmission of the infectious agent between individuals within clusters is an important mode of transmission, then the disease incidence in vaccinated clusters may be reduced by both direct and indirect protective effects of an efficacious vaccine. cRCTs can measure these combined effects, but only in special circumstances can they be designed and analyzed to elucidate the relative contributions of direct and indirect effects. In contrast, iRCTs only measure direct vaccination effects.^27^

#### D. Choosing a randomization strategy for a trial during an infectious disease emergency

When testing an investigational vaccine during an epidemic of an emerging infectious disease, it is urgent to establish an efficacy estimate before the epidemic gets too large (a factor at the beginning of an epidemic) or before cases become so rare that a trial is no longer feasible (a factor at the end stage of an epidemic). This urgency places a great premium on efficiency and speed of gaining a reliable result. iRCTs nearly always yield results that are easier to interpret and more precise than cRCTs involving the same number of participants.

The fact that iRCTs measure an investigational vaccine’s direct vaccine effects at the individual level, while cRCTs measure the combined direct plus indirect effects at the population level, may be arguments in favor of either design. By including indirect effects, cRCTs mimic the way that a vaccine would be used in public health programs; they may therefore measure a parameter – the population-wide reduction in disease from rollout of a vaccine – of direct interest to decision makers. However, this advantage may be offset by the fact that indirect effects are less readily transportable between settings. Also, regulatory decisions on the licensure of a vaccine are generally based on the estimates of the direct effects of the vaccine; herd effects are usually estimated in post-licensure studies. The indirect effects of a vaccine depend not only on the direct effects on vaccinated individuals, but on such factors as the incidence rate of the disease, the level of pre-existing immunity in the population, the contact network structure, the coverage of the vaccine and the phase of the epidemic (growing or declining), among others. Each of these factors will vary across populations that may consider using the investigational vaccine if it proves effective, and some of them (e.g. epidemic phase, immunity) may vary over short time periods within the same population. Dependence of indirect effects on setting, population, network structure or vaccine coverage level has been shown in models and studies of cholera, influenza, pneumococcal and other vaccines.^23,28–30^ Thus, a cRCT measures a parameter that is highly relevant to the time and place where it is measured, but which may be less relevant at a later time in the same population, or in a different population.

For example, the Ebola ça Suffit! trial^15^ was a cluster-randomized trial of an Ebola vaccine during the 2014–16 Ebola epidemic in which clusters were defined as those to whom an index cases was at high risk of transmitting the virus (the index case’s social contacts and their own social contacts) – called “ring” vaccination by analogy with the strategy used to control smallpox transmission in the end stages of the eradication program. This trial showed that in vaccinated individuals the investigational vaccine was highly protective (point estimate 100%). Effectively, the trial measured the combined direct and indirect effect of the investigational vaccine. The direct and indirect effects cannot be easily separated in this trial, and it is possible that the performance of the vaccine would be different if used in a different strategy to ring vaccination or in different settings within the same epidemic (i.e. urban vs. rural).^31^ Nonetheless, cRCTs do give a measure of the likely effect of the vaccine if deployed in a similar way in a similar setting.

The direct vaccine effect against infection measured in iRCTs is a more fundamental parameter from which, under plausible assumptions, indirect effects can be projected for various epidemic settings and investigational vaccine programs; working backwards from the indirect to the direct effects is much more complex and assumption-ridden. Thus, we posit that the most valuable parameter to estimate in most trials conducted during epidemics of emerging infectious diseases will be the direct effect of an investigational vaccine, which can only be directly measured in an iRCT.

Nonetheless, particular circumstances may weigh in favor of a cRCT. Recent work has shown that under certain circumstances, the difference in sample size requirements between a cRCT and an iRCT may be modest, because the larger effect measured by a cRCT partly offsets the design effect.^32^ Clearly, if a design is not feasible logistically or is unacceptable to the local population or health providers, it cannot be run effectively and produce valuable results. During the 2014–16 Ebola epidemic, it was argued that communities in the affected countries might be unwilling to accept individual randomization,^33^ particularly if, for example, different members of the same family might end up in different study arms. These claims require scrutiny, as what “the community” is willing to accept is both difficult to ascertain and depends on how the advantages and disadvantages of different trial designs are presented and discussed. For example, it was possible to launch an iRCT of an investigational vaccine in Liberia after extensive community engagement, albeit at a time when the epidemic was waning and there was less fear about Ebola.^34^ Nonetheless, if iRCT designs are found to be unacceptable and participants refuse or resist participation, this could threaten a trial’s social and scientific value—and thereby the fundamental ethical justification for conducting the trial. It might also undermine trust more generally and threaten the success of other epidemic control measures. cRCTs may also offer feasibility advantages over iRCTs, as all participants in one location are receiving the same intervention, eschewing the need for logistically more complex randomization procedures.

Another reason for choosing a cRCT design to improve a trial’s social and scientific value might (we hypothesize) arise in the setting of a declining epidemic, as was the case for all trials initiated to test investigational vaccines during the 2014–16 Ebola epidemic. If transmission has declined to low enough levels due to public health interventions, behavior change, depletion of susceptible hosts or other reasons, then an iRCT of an investigational vaccine that proved highly protective could reduce transmission effectively not only to vaccinated individuals, but to their unvaccinated contacts – i.e. would have an indirect effect (in both vaccinated and controls) even in the context of an iRCT. Thus, an iRCT that is designed to measure only direct effects might struggle to do so because the indirect effects of the trial investigational vaccine might reduce the number of cases in the control group. Such effects would depend upon the proportion of persons included in the trial who were at risk in the population from which trial participants were selected, and on the vaccinated:control ratio used in the trial. The extent to which such a problem, currently only hypothetical, might arise requires further analysis. While our understanding is still evolving, and there may be exceptions, the general principle remains that iRCTs provide a more efficient and reliable estimate of the direct vaccine effect than cRCTs, which we have argued is the fundamental measure of interest, rather than indirect vaccine effects. iRCTs are therefore likely to be the preferred design for evaluating experimental vaccines during epidemics of emerging infectious diseases.

### II. Trial population

#### Overview

Trial participants may either be selected from the general population or from a group at high risk of exposure to infection.

#### Features of a trial in the general population

Results of a trial performed in the general population may be the most appropriate if the vaccine is meant for widespread use in the general population, should it be shown to be efficacious in the trial. However, such trials will only be feasible if the incidence of the disease to be prevented by vaccination is sufficiently high to allow a trial of manageable size. Conducting a trial in the general population tends to equalize everyone’s chances of participation. While, in practice, the socially or politically powerful may have better access to trial locations or be permitted to enter trials of promising interventions ahead of the less powerful, recruiting the general population still promotes more equal opportunity for access to an investigational product that many might want to receive during an epidemic.

#### C. Features of a trial in a population at high risk of exposure

If the disease is relatively uncommon, detecting a beneficial effect of the investigational vaccine in the general population with adequate statistical power might require a prohibitively large trial. A trial population comprising individuals at high risk of exposure to infection, such as sero-discordant couples for a sexually transmitted infection^35^ or health care workers for a disease transmitted by direct contact,^36^ is likely to have greater statistical efficiency. Moreover, at-risk populations are more likely to benefit if the vaccine under study proves beneficial, and they are likely to become a priority group for any future vaccination program. Targeting an at-risk population can thereby enhance a trial’s social and scientific value. Given that the investigational vaccine is being tested for a putative beneficial effect that depends on exposure to the pathogen, while certain adverse effects may occur in both unexposed and exposed vaccine recipients, it is likely that the expected benefit would be larger per participant in a highly exposed population.

Targeting an at-risk population can also promote fair participant selection, depending on how fairness is defined. When a population is at increased risk of infection because they are undertaking important public health work, such as burying the dead, or caring for patients or ill family members, targeting this population promotes fairness as reciprocity. At the same time, focusing a trial on certain at-risk populations, such as front-line health workers, may invite charges of conducting the trial in those of relatively high social standing and undermining efforts to equalize access to the investigational vaccine more generally. Ethical debate about which notion of fairness is most convincing in this context is ongoing.^37,38^

#### D. Choosing a population for a trial during an infectious disease emergency

Efforts to enhance a trial’s risk-benefit profile may lead to performing a trial in a group that is especially likely to benefit if the investigational vaccine proves efficacious, such as those with occupational or familial/household exposure to the infection. This approach can also enhance a trial’s social and scientific value by reducing the sample size required to conduct an adequate trial, given the increased risk of infection. Similarly, efforts to enhance a trial’s risk-benefit profile may favor excluding those who are most at risk from possible adverse effects of the investigational vaccine; these may, in various situations, include children, pregnant women and the fetuses they carry, or individuals with particular medical conditions, such as immune deficiencies. However, if such individuals would be in the eventual target population for a vaccination program, there are compelling arguments for including them in a trial, which would include an assessment of safety in such groups.

Complexities ensue when these considerations come into conflict. Those who may be at greatest risk from an investigational vaccine – who might be excluded from trials – may be also those who stand the most to gain if it is beneficial. An example is pregnant women for investigational vaccines against Zika virus infection; concern about adverse events on the fetus might argue for excluding such women, but they are clearly also the ones who could benefit the most if the investigational vaccine proves effective. Excluding women of childbearing age (who might be pregnant) or pregnant women can make a trial’s risk-benefit profile less favorable, depriving the group of potential clinical benefits from participating in the trial and also thereby reducing the trial’s scientific and social value because data on a key target population for the vaccine are not collected. Importantly, these considerations are also relevant for judging whether a trial’s participant selection is fair, as compelling reasons are required for excluding entire population groups. In the case of women of childbearing age and pregnant women, a systematic precautionary approach has led to the unfair exclusion of both groups over decades.^39,40^ The default should therefore be to include women of childbearing age and pregnant women—as well as other so-called vulnerable groups—in investigational vaccine trials during epidemics of emerging infectious diseases, provided that the risks and potential benefits of participating in the given trial are acceptable.^12,41^

For infections that are naturally immunizing, investigators might choose to restrict enrollment to those who have not previously been infected to ensure that trial participants are truly at risk of becoming infected – this is especially relevant when selecting persons thought to be highly exposed. However, selecting participants who both have risk factors for infection and are uninfected at enrolment – as in a study in sero-discordant couples – can lead to additional issues. First, it complicates the study conduct as all potential participants must be tested for evidence of prior infection. Second, individuals who have remained uninfected despite many opportunities for exposure may be more resistant to infection (or may have lower-risk exposures) than is typical in the population.^42^ Sero-discordant couples, for example, may tend to be those that practice safer sex or in which the infected partner is relatively less infectious than in other couples; likewise, health-care workers who remain uninfected despite intense exposure may practice better personal protection than is typical. Failure to account for such factors may lead to overly optimistic estimates of statistical power through over-estimating the likely infection rate during the trial;^42^ moreover, if the vaccine is differently effective in different populations, then the effect estimate from such a trial may be biased compared to what it would be for the general population.^24,25,43^ Infections, such as dengue, which confer more complex types of immunity, including enhancement, require special consideration outside the scope of this paper.^44^

Finally, feasibility considerations can influence who is selected for inclusion in the trial population during epidemics of emerging infectious diseases. For example, if the supply of the investigational vaccine is limited and difficult to manufacture to scale before the window of opportunity for conducting a trial closes, this can justify increasing the efficiency of the design by targeting at-risk populations.

### III. Comparator intervention

#### Overview

In a randomized trial, investigators must choose the intervention that the control arm will receive. In general, those in the control arm may receive a placebo, or another vaccine which is unlikely to affect the disease under study. Ideally, the control intervention should be identical in appearance to the investigational vaccine, such that both the investigators and the participants will be unaware of who is in which arm (double-blind design). In practice this is often difficult to achieve and blindness is maintained by ensuring that those who administer the investigational vaccine or control play no part in assessing endpoints of interest in the trial and, to the extent possible, that participants are unaware of which intervention they receive.^27^ Alternatively, control participants may receive nothing or receive the investigational vaccine at a later point during the trial, where the vaccine’s effects are compared between the different temporal stages.

#### B. Features of placebo comparator

Use of a placebo control and blinding enhances a trial’s social and scientific value: if no one knows who is receiving the investigational vaccine or the control, there is no potential for bias in the assessment of trial endpoints between the two groups. In non-blinded trials, bias can arise in intervention allocation, if, for example, the investigators manipulated the randomization and knowingly put the more or less vulnerable participants in the investigational vaccine arm, or if participants change their behavior if they know they did or did not receive the investigational vaccine.

#### C. Features of active comparator

The justification for using an active control – most commonly a proven effective vaccine against another infection – is that some benefit is being provided to those in the control group, albeit not with respect to the disease under study, and this may enhance a trial’s risk-benefit profile. This is especially the case when the active control can be administered in such a way as to allow for double-blinding and the trial results are unlikely to be confounded by the comparator vaccine, such that the trial’s social and scientific value rests ensured. For example, in the Phase 3 trial of the RTS,S/AS01 malaria vaccine, the rabies vaccine was used as an active control.^45^

#### D. Features of delayed-administration comparator

Comparison against delayed administration, without use of a comparator vaccine or placebo, is a way for the investigators to ensure every participant is offered the investigational vaccine at some point. In situations where the investigational vaccine is thought likely to be efficacious, this can enhance the risk-benefit profile of the trial. The major disadvantage of this approach is that individuals clearly know if they are in the vaccine or control group and this may lead, for example, to differential behavior changes in the two groups, affecting their risk of disease independently of any biological protective effect of the vaccine. Another disadvantage in cRCTs, if a comparator vaccine is not used contemporaneously in the control group, is that in the vaccine arms it is possible to clearly identify those who were vaccinated and those who refused vaccination, whereas if no intervention is offered in the control arm at the start of the trial it may be difficult or impossible to identify those who would have been vaccinated had they been offered the vaccine. As acceptors and refusers may be at differential risk of disease, independently of vaccination, the only unbiased comparisons that are possible are to compare disease rates among all those in the vaccine clusters and those in the control clusters, which may result in an under-estimation of the vaccine’s effect if substantial numbers refuse vaccination.

Of course, in a trial with the use of a comparator vaccine or placebo in the control arm, it is still possible to offer all those in the control arm the experimental vaccine after it is shown to be efficacious in the trial. However, any delay in providing the vaccine requires the control group to forego potential benefits.^46^ A disadvantage is that providing the experimental vaccine in the control arm will prevent assessment of the longer-term efficacy of the vaccine and may obscure the observation of adverse effects that may arise in the longer term.

#### E. Choosing a comparator during an infectious disease emergency

Compared to both an active control and a delayed-intervention control, use of a placebo control is likely to maximize a trials’ scientific value because of the opportunity for blinding and the lack of concern that the incidence of the disease under study in the control group may be affected in some way by the placebo intervention; all else equal, it is the preferred comparator. Use of an active control can complicate the assessment of safety for the investigational vaccine as only comparative safety between the two interventions will be measured. Even in situations where it is judged in advance that an active control will not have either protective or adverse effects that could be confounded with effects of the experimental vaccine, there is the risk that this judgment could be wrong.^47^ However, using delayed administration as a comparison can also complicate the long-term assessment of efficacy, as after both arms receive the intervention there is no comparison group that has not received the vaccine. Thus, to enhance a trial’s social and scientific value, a placebo control should generally be chosen. However, if an active control does not undermine a trial’s scientific value, it should be used to enhance the trial’s risk-benefit profile.

During an epidemic, political leaders, community representatives or others can have a strong preference for ensuring that all trial participants have access to the investigational vaccine - especially if prior evidence suggests it is likely to be efficacious (e.g. protection shown in non-human primate studies and high immunogenicity), the disease is serious, the burden of disease is high, and available preventative or therapeutic measures are limited. If this preference makes using a placebo control unfeasible, comparing the investigational vaccine to delayed administration can be justified provided the social and scientific value of the study is not undermined.^46^

### IV. Trial implementation

#### Overview

In the simplest experimental trial design, all trial participants are enrolled into the trial concurrently and followed for the same period of time, known as a “parallel” design trial. However, sometimes it is not feasible to enroll all participants at once and a “stepped” rollout is used, wherein entry to the trial is phased over time. While some degree of stepped rollout is in almost all trials (as all participants cannot be enrolled and vaccinated on the same day), it has been used as a deliberate design choice in cRCTs, in the form of the “stepped wedge” design. This design was first used to evaluate the introduction of the hepatitis B vaccine in The Gambia,^48^ and it entails a phased introduction of the vaccine over time. In a stepped wedge cRCT, the order of introduction to the various clusters of groups is randomized, and the disease incidence is then compared in successive time intervals, between those clusters in which the intervention has already been rolled out and those in which it has not yet been rolled out. The appeal of this design is that by the end of the study all clusters have received the investigational vaccine. A stepped wedge design is typically used to evaluate previously tested interventions in new settings. It has never been used to evaluate an unlicensed investigational vaccine, although it was proposed for the evaluation of an investigational vaccine against Ebola in Sierra Leone among frontline health workers.^49^

#### B. Features of parallel rollout

When sufficient supplies of the investigational vaccine and control interventions (if any) are available at the start of a trial; when the geographic area for the trial is clearly identified (and anticipated to have continuing disease transmission throughout the trial); and when logistics permit large-scale rollout simultaneously to the entire trial population, starting the trial at a similar time in all participants will minimize the time required to obtain a result. If these criteria are met, there is no reason to delay the enrolment of participants into the trial.

#### C. Features of stepped rollout

Deliberate rollout of the investigational vaccine over a period of time may be implemented due to limited supply of the investigational vaccine or limited capacity to implement the intervention in many locations simultaneously. In this setting, a stepped rollout can permit prompt commencement of the trial, without waiting until supplies or logistics make rollout everywhere feasible, and can permit a more rapid trial. Additionally, when supplies of a potentially promising investigational vaccine – or the personnel and other resources to deliver it – are scarce, one comparatively fair way to allocate it is to choose randomly who receives it, or if the supply is being augmented during the trial, to choose randomly who receives it first. Low and variable incidence of a disease may make a classic parallel trial rollout unfeasible because the trial can gain sufficient power only by targeting at-risk populations and randomizing individuals or clusters in the vicinity of known or predicted cases.^15,49,50^ Such trials, which follow the cases in real time, are rolled out necessarily in a stepped fashion.

#### D. Choosing a rollout strategy in an infectious disease emergency

The stepped-wedge design – a cRCT using stepped rollout – has been used as a way to roll out an intervention that is likely to be effective so that all eventually receive the intervention, but it is introduced in such a way that assessment of effectiveness is possible. The design might also be used if the intervention could not be provided everywhere simultaneously, due, for example to a supply shortage, while evaluating the effects of the intervention as it is introduced over time.^51^ Such a design was considered for this reason during the Ebola outbreak in 2014–6, especially when supplies of investigational vaccines were limited. However, an analysis comparing this approach to a parallel iRCT showed that the stepped-wedge cRCT would suffer from inadequate power given the variable and unpredictable incidence, so the parallel iRCT was preferred from the perspective of social and scientific value.^19^ To combine the advantages of stepped rollout with individual randomization, it was proposed to conduct an adaptively designed iRCT, using an individual randomization strategy but rolling out the investigational vaccine supply as it became available (“stepped rollout”).^49^ It has been argued that such a design would capture many of the benefits of a stepped-wedge trial while also achieving the greater efficiency of individual randomization.^52^

The Ebola experience shows that the conditions favoring stepped rollout – limited supplies of the investigational or comparator vaccine, limited logistical abilities to conduct a trial (especially without hindering other response efforts), and variable incidence that supports conducting the trial in areas of known incidence – can readily occur in an emergency setting. Indeed, this was an important part of the justification for the design that was ultimately used with success to test an Ebola vaccine in the Ebola ça Suffit! Trial, which used a form of stepped rollout by enrolling participants from the contacts and contacts of contacts when cases arose. Notably, some conditions (limited supplies) were particularly acute at the start of the epidemic, while another (variable incidence) was particularly pressing at the end of the epidemic. For this choice, there is no general preference for one option over another: details of the particular emergency (which may change rapidly as it progresses) will have to inform the rollout option chosen in order to ensure or enhance the social and scientific value of the trial.

## Conclusion

We have assumed a setting in which no licensed vaccine is available and have focused on experimental or randomized designs for testing an investigational vaccine during epidemics of emerging infectious diseases. We acknowledge that the considerations will change if expanded to other contexts in which more interventions are available or if there are settings in which randomization is not possible.^11^

We have limited our scope to decisions we believe could most delay the start of a trial for an emerging infectious disease with no proven vaccine. A closely overlapping set of decisions has been incorporated into an online interactive decision tool (InterVax-Tool: http://vaxeval.com/).^53^ We did not consider subsequent design choices, such as sample size, trial duration, or the choice of trial endpoint^53,54^ (e.g., infection vs. symptomatic infection), as these will be dependent on the particular pathogen in question. Similarly, there are many important ethical considerations such as informed consent and community acceptance that can tip the balance when choosing between several possible designs. For example, if two designs are acceptable given their social and scientific value, risk-benefit profile and participant selection, but they differ in how the designs are understood by participants, then the design that is better understood should be chosen in order to promote informed consent. Finally, we did not address regulatory issues beyond the scientific considerations that often underpin decisions about vaccine licensure. Because licensure is a key goal of any vaccine trial, regulatory considerations are essential for choosing between trial designs during epidemics of emerging infectious diseases. However, a thorough comparative discussion of different regulatory regimes and their implications for trial choices goes beyond the scope of this paper.

By no means have we detailed all decisions that must be made regarding trial design. However, through a discussion of four key elements that must be considered when choosing trial designs, we hope to have highlighted the range of options of designs that should be considered during future outbreaks of emerging infectious diseases.

## Acknowledgments

We thank Ethox Centre at Oxford University for supporting the workshop that led to the writing of this paper, which draws content from some of the materials distributed in advance of that meeting by the organizers (RK NE AR ML) and from the presentations of AR and PS there.

